# Transcriptomic heterogeneity of *Sox2*-expressing pituitary cells

**DOI:** 10.1101/2021.12.10.472137

**Authors:** Patrick A. Fletcher, Rafael M. Prévide, Kosara Smiljanic, Arthur Sherman, Steven L. Coon, Stanko S. Stojilkovic

## Abstract

The mammalian pituitary gland is a complex organ consisting of hormone-producing cells (HPC), nonhormonal folliculostellate cells (FSC) and pituicytes, vascular pericytes and endothelial cells, and putative *Sox2*-expressing stem cells. Here, we used scRNAseq analysis of adult female rat pituitary cells to study the heterogeneity of pituitary cells with a focus on evaluating the transcriptomic profile of the *Sox2*-expressing population. Samples containing whole pituitary and separated anterior and posterior lobe cells allowed the identification of all expected pituitary resident cell types and lobe-specific subpopulations of vascular cells. *Sox2* was expressed uniformly in all FSC, pituicytes, and a fraction of HPC. FSC comprised two subclusters; FSC1 contained more cells but expressed less genetic diversity compared to FSC2. The latter contained proliferative cells, expressed genes consistent with stem cell niche formation, including tight junctions, and shared genes with HPC. The FSC2 transcriptome profile was also consistent with the activity of pathways regulating cell proliferation and stem cell pluripotency, including the Hippo and Wnt pathways. The expression of other stem cell marker genes was common for FSC and pituicytes (*Sox9, Cd9, Hes1, Vim, S100b*) or cell type-specific (FSC: *Prop1, Prrx1, Pitx1, Pitx2, Lhx3;* pituicytes: *Fgf10, Tbx3, Lhx2, Nkx2-1, Rax*). FSC and pituicytes also expressed other astroglial marker genes, some common and other distinct, consistent with their identities as astroglial cells of the pituitary. These data suggest functional heterogeneity of FSC, with a larger fraction representing classical FSC, and a smaller fraction containing active stem-like cells and HPC-committed progenitors.

## 1 INTRODUCTION

The pituitary gland consists of two parts: the anterior pituitary (or adenohypophysis) composed of the anterior and intermediate lobes, and the posterior pituitary (or neurohypophysis) composed of the posterior lobe and the pituitary stalk or infundibulum, which connects the posterior lobe to the hypothalamus. The gland can be considered to contain three groups of resident cells: hormone producing corticotrophs, gonadotrophs, thyrotroph, somatotrophs, and lactotrophs located in the anterior lobe, and melanotrophs located in the intermediate lobe; nonhormonal folliculostellate cells (FSC) in the anterior pituitary and pituicytes of the posterior pituitary; and endothelial cells (EC) and pericytes from the vascular networks of the anterior and posterior pituitary. Hormone producing cells (HPC) are well characterized (Stojilkovic et al., 2010), but their homogeneity vs. heterogeneity/plasticity is still questioned (Childs et al., 2020). The nonhormonal FSC were first described almost 70 ago (Rinehart & Farquhar, 1953) and characterized as stellate cells forming anterior pituitary spanning networks (Fauquier et al., 2001; Vila-Porcile, 1972). Significant progress in understanding their structure, diverse functions, and genetic profiles has been achieved, predominantly in work by Yukio Kato, Kinji Inoue, and colleagues, aided by the development of an S100b-GFP transgenic rat model (Devnath & Inoue, 2008; Itakura et al., 2007; Kato et al., 2021). Pituicytes are posterior pituitary glial cells (Verkhratsky & Nedergaard, 2018), with similar genetic and functional profiles as tanycytes (Clasadonte & Prevot, 2018). Based on histological and physiological features, FSC (Allaerts et al., 1996; Horiguchi et al., 2010) and pituicytes (Virard et al., 2008) are considered as a heterogeneous subpopulation. Pericytes and EC are the least characterized pituitary cells, but the relevance of hypothalamic-vascular-pituitary unit other than blood supplies was recently raised (Le Tissier et al., 2017). However, genetic heterogeneity within hormonal and nonhormonal cells is difficult to characterize without transcriptomic measurement at single cell resolution.

The presence of postnatal pituitary stem cells was also proposed to account for pituitary cell proliferation in response to a variety of factors, such as pituitary growth during prepubertal period (McNicol & Carbajo-Perez, 1999), adult pituitary cell maintenance (Langlais et al., 2013), osmotic stimulation (Virard et al., 2008), injury (Fu et al., 2012), and after gonadectomy, adrenalectomy (Rizzoti et al., 2013), and estrogen treatment (Mitsui et al., 2013). These cells express genes characteristic of stem/progenitor cells, including *Sox2*, and have proliferative capacity (J. Chen et al., 2005; Fauquier et al., 2008). They are thought to reside in two types of stem cell niches: the marginal layer of cells lining Rathke’s cleft between the anterior and intermediate lobes and dense clusters dispersed throughout the gland parenchyma (S. Yoshida et al., 2009). These cells have also been reported to express other stem cell genes shown to contribute to embryonic development of resident cells, such as *Cdh1, Cd9, Lhx3, Prop1, Prrx1, Vim, Sox9*, and *S100b* (Gleiberman et al., 2008; Horiguchi et al., 2018; Rizzoti, 2010; Vankelecom & Chen, 2014). It has also been suggested that FSC act as stem cells in the anterior pituitary gland (Horvath & Kovacs, 2002; Inoue et al., 2002; Vankelecom, 2007), similarly to brain astrocytes (Osuna et al., 2012). In the posterior lobe, pituicytes are continuously replaced by proliferation and differentiation of stem-like cells (Virard et al., 2006). *Rax* appears to be a gene of interest for defining pituicytes generating cells (Pak et al., 2014). The possibility that neurohypophysial progenitor cells originate from stem cells in the marginal zone has also been considered (Miyata, 2017). However, the experimental protocols used in these studies did not allow the characterization of the genetic profile of pituitary stem cells.

Single cell RNA sequencing (scRNAseq) provides the capability to study pituitary cell transcriptome profiles and heterogeneity at single cell resolution, making it an important tool for identifying pituitary stem cell cluster(s), for comparing FSC with known stem cell expression profiles, and for advancing the understanding of the potential roles of EC and pericytes in pituitary cell functions. Initial scRNAseq studies were helpful in the global characterization of HPC in male mice (Cheung et al., 2018), male and female rats (Fletcher et al., 2019), and human fetal pituitaries (Zhang et al., 2020), as well as pituicytes from male mice posterior pituitary (Q. Chen et al., 2020). *Sox2*-positive cells have been detected in pituitary cells in these studies, but contradictory conclusions have been drawn about the cell types expressing this gene. We and others reported that *Sox2*-expressing cells expressed genes consistent with FSC identity (Fletcher et al., 2019; Vennekens et al., 2021), in agreement with immunohistochemical studies (Kato et al., 2021). Others have defined *Sox2*-positive cells uniformly as stem cells (Cheung & Camper, 2020; Cheung et al., 2018; Lopez et al., 2021; Mayran et al., 2019; Zhang et al., 2020), effectively renaming FSC. Moreover, comparative transcriptome profiling of FSC, pituicytes, HPC, pericytes, and EC has not yet been reported.

To help gain insights into the transcriptomic profile of *Sox2*-expressing cells and their relationship to HPC, nonhormonal cells, and vascular cells, we present here scRNAseq data on cells from freshly dispersed whole pituitary glands of adult random cycling female rats. While we did not find evidence of multihormonal/multireceptor types of HPC, we did observe lobe-specific clusters of vascular cells, posterior lobe pituicytes, and two subclusters of anterior lobe FSC. We show that *Sox2* and *Sox9* are not only the marker genes of stem-like cells in the pituitary gland but are also present in “classical” FSC and pituicytes, that several reported stem marker genes also shared expression in vascular cells or HPC, and that others were FSC or pituicyte-specific. Our result support the hypothesis that a small subset of cells in the pituitary FSC network have stem cell niche and progenitor activity involved in basal endocrine cell renewal in the adult gland. Finally, we provide support for the common identity of FSC and pituicytes as pituitary astroglial cells.

## 2 MATERIALS AND METHODS

### 2.1 Animals

Sprague Dawley rats were obtained from Taconic Farms (Germantown, NY) and housed for two weeks under constant temperature and humidity conditions, with light on between 6 and 20 h. The experiments were performed with 75-day-old random cycling females. After decapitation and removal of the brain, the whole pituitaries were collected, or the posterior/intermediate and anterior lobes were separated prior to collection; glands were used for RNA extraction or cell dispersion as described below. All experimental procedures were approved by the National Institute of Child Health and Human Development, Animal Care and Use Committee (Animal Protocol 19-041).

### 2.2 Dispersion of pituitary cells

Dissociation of cells was performed using 30 whole pituitary glands or 30 separated anterior and intermediate/posterior pituitary lobes and trypsin/EDTA based method. Briefly, the glands were quickly collected and kept in ice-cold 199 Hanks media. The tissue was then washed with ice-cold phosphate buffer saline (PBS) supplemented with 0.3% bovine serum albumin fraction V (MP Biomedical, Solon, OH) and 1.26 mM CaCl_2_ (Quality Biological, Gaithersburg, MD). All collected tissue was chopped in 0.5 × 0.5 mm pieces and incubated in a water bath shaker for 15 min at 50 rpm, in the presence of a trypsin solution (4 mg/ml; Sigma, St Louis, MO), which was then replaced with a solution containing 2 mg/ml trypsin inhibitor (Sigma) for 5 min. The latter was removed, and 2 mM EDTA (Corning, Manassas, VA) solution was added over 5 min and then replaced with a 1 mM EDTA solution over 10 min with constant shaking. The volume of media was proportional to the weight of the tissue. After these steps, tissue was mechanically dispersed, centrifuged, the cells were counted and immediately used for scRNAseq, RNA isolation, or plated for immunocytochemistry.

### 2.3 Quantitative RT-PCR

RNA was extracted from individual anterior and posterior pituitary glands, and anterior pituitary cells 30 min after cell dispersion, using a RNeasy Plus Mini Kit (Qiagen, Valencia, CA). RNA was reverse transcribed with a Transcriptor First Strand cDNA Synthesis Kit (Roche Applied Sciences, Indianapolis, IN). Quantitative RT-PCR was performed using Applied Biosystems pre-designed TaqMan Gene Expression Assays for rats and TaqMan® Fast Advanced Master Mix. PCR was performed in the QuantStudio 6 Flex Real-Time System (Applied Biosystems, Waltham, MA). Target gene expression levels were determined by the comparative 2_∧_-(delta C(T)) quantification method using *Gapdh* as the reference gene, which was previously established to be a suitable reference gene for the rat anterior pituitary tissue (Janjic et al., 2019). Applied Biosystems predesigned TaqMan Gene Expression Assays were used: *Aldh1a1* (Rn00755484_m1), *Aldh1l1* (Rn00674034_m1), *Aqp4* (Rn01401327_s1), *Edn3* (Rn01755284_m1), *Fibin* (Rn01514386_s1), *Gapdh* (Rn01462662_g1), *Gfap* (Rn01253033_m1), *Gstm2* (Rn00598597_m1), *Maob* (Rn00566203_m1), *Nes* (Rn01455599_g1), *Penk* (Rn00567566_m1), *S100a1* (Rn01458753_m1), *S100a6* (Rn00821474_g1), *S100a10* (Rn06378613_s1), *S100a11* (Rn01409258_g1), *S100a13* (Rn01769833_m1), *S100a16* (Rn01458849_g1), *S100b* (Rn04219408_m1), *Slc1a2* (Rn00691548_m1), *Slc1a3* (Rn01402419_g1), *Sox2* (Rn01286286_g1), *Sox9* (Rn01751070_m1), *Sult1a1* (Rn01510633_m1), and *Vim* (Rn00667825_m1).

### 2.4 Immunohistochemistry

For immunocytochemical analysis, 50,000/well freshly dispersed anterior pituitary cells were plated on poly-L-lysine-coated 8-well multitest slides (MP Biomedicals, Aurora, OH), bathed in Earle’s medium 199 supplemented with 2.2 g/L sodium bicarbonate, 10% heat-inactivated horse serum, 100 units/ml penicillin, and 100 μg/ml streptomycin. Following overnight incubation, cells were washed with PBS two times, fixed with 4% formaldehyde solution (Thermo Scientific, Rockford, IL, USA) for 10 min at room temperature and incubated either with anti-S100A (1:400; GenTex, Irvine, CA) or anti-S100B (1:500, Novus Biologicals, Centennial, CO) antibodies overnight at 4 °C, followed by incubation with appropriate secondary antibody (Alexa Fluor 488 donkey anti-mouse or Alexa Fluor 488 goat anti-rabbit; 1:1000) for 30 min at room temperature. For double immunolabeling, cells were incubated with both anti-S100A and anti-S100B antibodies. All antibodies were diluted in staining PBS solution containing 0.2% saponin and 0.5% BSA. Every step of immunostaining protocol was followed by washing cells three times with PBS. To confirm the specificity of the reaction, negative controls were generated by omitting the primary antibodies during the immunohistochemical procedure (data not shown). Cells were mounted with Fluoromount-G (Invitrogen, Carlsbad, CA) containing 4′,6-diamidino-2-phenylindole (DAPI) as a nuclear counterstain. All images were acquired on an inverted confocal laser-scanning microscope (LSM 780; Carl Zeiss GmbH, Jena, Germany), using 63x Oil objective. Micrographs were sized, and their brightness and contrast levels adjusted in Fiji. Cells were counted on 12 tile-scan images (3×3).

### 2.5 Single cell RNA sequencing

#### 2.5.1 Library preparation, sequencing, and transcript counting

Dispersed cells from two cell preparations were used, one whole-pituitary and the other with anterior and posterior pituitaries separated. The whole pituitary sample was run in duplicate and the anterior and posterior samples each run in separate lanes, for a total of four lanes on the 10X Genomics chromium controller loaded at 9000 cells per lane according to manufacturer instructions. Resulting libraries were sequenced on an Illumina HiSeq 2500, and the Cellranger pipeline (10X Genomics) was used for mapping reads and counting transcripts. A custom reference genome was used as previously described (Fletcher et al., 2019), based on the Ensembl Rnor6.0 genome, extended up to 6kb upstream or until the next feature was encountered. The four samples were aggregated with CellRanger aggr, resulting in a total of 32455 droplets, with a mean of 20950 reads, a median of 1803 unique genes, and a median of 5693 UMI per droplet. The data analyzed in this manuscript is publicly available at Genome Expression Omnibus accession number GSE184319 (https://www.ncbi.nlm.nih.gov/geo/query/acc.cgi?acc=GSE184319).

#### 2.5.2 Dimensionality reduction, visualization, and clustering

For visualization and reporting of expression levels, transcript counts were normalized to total cell library size and log transformed with pseudocount of 1. Dimensionality reduction was performed in Seurat v4.1 using SCTransform normalization of raw counts with 2000 variable genes (glmGamPoi method), followed with PCA, retaining 50 components (Hao et al., 2021). Distances between cells in the resulting PC space were measured using correlation distance and used for subsequent k-nearest neighbor (kNN) identification (knnsearch in Matlab R2021b) and UMAP embedding (umap-learn version 0.5.1) (Mcinnes et al., 2018). Unsupervised clustering of the graph generated by UMAP was performed using the Leiden algorithm (leidenalg version 0.8.1) (Traag et al., 2019). Violin plots were made using ViolinPlot (Bechtold, 2016), and all other plots were generated using Matlab 2021b (The MathWorks, Inc.).

#### 2.5.3 Cell filtering

The transcriptomes of cell types of the pituitary are heterogeneous, so cell quality control was performed in a cluster-based manner. First, unsupervised clustering was performed (resolution=1) and the distributions of cell quality metrics were inspected per cluster. Clusters composed entirely of droplets with low number of genes and high fraction of counts attributed to mitochondrial transcripts (fracMT) were identified and removed. In the remaining clusters, thresholds for high fracMT and low total gene count were defined for each cluster as the median ± 3 median absolute deviations, respectively. A global lower limit of 750 genes and upper limit of 30% mitochondrial fraction was used in cases where the adaptive thresholds exceeded these values. This procedure filtered 14828 droplets. Also removed were a small cluster of erythrocytes (164 cells), and two small clusters of unidentified cells from the posterior lobe sample, which we deemed too few to warrant further analysis (72 cells total).

#### 2.5.4 Cell classification and identification of ambiguous cells

To address the possibility of multiplet cells or putative multihormonal cells, we used a cell-wise classification strategy based on cell type marker genes. This approach allowed for the identification and analysis of cells expressing signatures of more than one cell type. Cell clusters remaining after outlier removal could be readily identified for each expected pituitary cell type based on canonical marker genes for expected pituitary cell types. We used this initial classification to find genes with dominant expression in each cell type cluster, focusing on those with expression in at most 5-10% of cells of all other types. This refined set of markers was then used to compute cell type scores for each cell type, per cell, using a Matlab implementation equivalent to Seurat’s AddModuleScore. Pairwise scatterplots of the marker scores were inspected to determine thresholds above which cells were assigned to a cell type. Cells that were uniquely classified were those that had scores above threshold in only one cell type score, appearing near the x- or y-axis only. Ambiguously classified cells, which were above threshold in more than one cell type marker score, appeared as clusters of cells near the diagonal (1492 cells). A small number of cells could not be classified, having no cell type score above threshold (23 cells). These cells were present primarily among clusters of uniquely classified cells, so we attempted to assign their cell type using kNN classification based on the most common identity of their neighbors. To finalize the cell filtering, ambiguous cells and remaining unclassified cells were excluded from further analysis, leaving 15876 cells. The remaining unambiguously classified cell types corresponded very closely to UMAP clusters, confirming that cell-wise classification was successful.

#### 2.5.5 Ambient RNA removal with SoupX

We found significant contamination of droplets by ambient RNA, particularly for hormone genes such as *Prl, Gh1, Pomc*, and *Cga*. We quantified the contribution of genes to this ambient RNA with SoupX (Young & Behjati, 2020) and used SoupX adjusted counts for differential expression analysis and visualization. Cell type identities were used as input clustering for SoupX and used the automatic estimation mode on each sample. The estimated global contamination fractions (rho values) were in 0.01-0.02 for all four samples, but we found that the contamination fraction needed to be set higher to remove hormone gene contamination; we used the upper value of rhoFWHM, which ranged from 0.039 - 0.047.

#### 2.5.6 Differential expression analysis

Differential gene expression was done using a pairwise comparison approach on SoupX corrected counts to identify upregulated genes in cell types of interest relative to other cell types as previously described (Fletcher et al., 2019). We used the ANOVA-like Kruskal-Wallis test (the extension of the Wilcoxon ranksum test to more than two groups) to identify whether a gene showed a difference in expression among cell types, followed by multiple comparisons to compute Bonferroni-adjusted p-values for each pairwise comparison (Matlab’s kruskalwallis and multcompare functions). The Kruskal-Wallis and pairwise p-values per gene were then corrected for multiple comparisons using the Benjamini-Hochberg false discovery rate (Groppe, 2021). Unless otherwise stated, cell type-dominant genes were defined as genes with expression in at least 15% of cells and with a significant upregulation (adjusted p-value < 1e-6) of at least 2-fold in normalized counts and a difference in proportion of expressing cells of at least +10%, relative to other cell types. Highly specific marker genes for a cell type were identified as above with the additional constraint that all other cell types had fewer than 5% of cells expressing the gene.

#### 2.5.7 Gene Ontology and pathway enrichment analysis

Gene Ontology (GO) and pathway enrichment was performed using the python implementation of gProfiler (Raudvere et al., 2019), with query gene sets for cell types of interest determined from differential gene expression analysis. We used unordered mode and the default p-value computation method (gSCS) with a threshold of p=0.05. For GO terms, we took advantage of the GO tree structure to remove redundant terms in from the set of enriched terms. Redundant terms could be identified as larger, more generic terms with a set of genes enriched that was identical to one of its descendent terms, following only “is_a” relations (Jantzen et al., 2011). To aid in interpretation of results from multiple queries, enriched terms were clustered using a gene-overlap approach. The sets of all unique terms and genes were identified across all queries. A term-gene matrix was used to represent, for each term, the number of queries for which each gene was enriched. The Jaccard distance metric was then used for UMAP embedding of the terms (rows of the term-gene matrix), and subsequent clustering of the UMAP graph was performed using leidenalg.

#### 2.5.8 Ligand-receptor interactions using CellChatDB

To examine the possibility of ligand-receptor interactions between cell types, we used the approach of CellChat (Jin et al., 2021) with some modifications. We first converted the gene names in the CellChat database of interactions from mouse to their rat orthologs using gProfiler’s orth tool. Next, we identified the set of genes identified from the database that were expressed in our dataset in at least 5% of cells of any cell type, which resulted in 459 genes representing 673 interactions. We normalized the expression of each gene to the maximum expression per gene, then computed the mean expression per gene, per cell type. The interaction strength for an interaction involving ligand L and receptor R was computed as sqrt(L*R), for every pairwise combination of cell types. If L or R were complexes, their expression value was the geometric mean of the expression of each member of the complex. We did not consider co-factors or activators and inhibitors, so as not to make any assumptions about the relationship between gene expression level, ligand and receptor protein levels, and ligand-receptor interaction affinities. Instead, we interpret the interaction strength as a simple potential for interaction due solely to gene expression levels. P-values were computed as is done in CellChat. A null distribution of interaction strengths was generated from randomized expression profiles via shuffling of cell labels, repeated 10000 times, and the observed interaction strength’s rank into this distribution was taken as the p-value.

## 3 RESULTS

### 3.1 The resident cell types of the pituitary gland

We performed single cell RNA sequencing on dispersed pituitary cells from 75-day-old random cycling female rats in two cell preparations, one with 30 whole pituitary glands and the other with 30 separated anterior and intermediate/posterior pituitary lobes. A total of 32455 droplets were recovered from four lanes of the 10X Chromium Controller platform (10X Genomics); two replicates of the whole pituitary sample, and one lane each for anterior lobe and posterior lobe samples. Dimensionality reduction for downstream analysis was done by first normalizing transcript counts with SCTransform from Seurat v4.1 (Hao et al., 2021), then performing principal component analysis followed by UMAP embedding (Mcinnes et al., 2018) and unsupervised graph-based clustering with the Leiden algorithm (Traag et al., 2019). Low quality droplets were identified based on low number of detected genes and high fraction of mitochondrial counts, using adaptive cluster-specific thresholds based on the median ± 3 median absolute deviations, filtering 14828 droplets (see Methods). Significant levels of ambient RNA were observed, most prominently for hormone genes including *Prl, Gh1, Pomc*, and *Cga*, so we applied SoupX to remove contaminating transcript counts for all further analysis (Young & Behjati, 2020).

Initial examination of expression patterns of known marker genes showed that cell clusters corresponding to expected resident pituitary cell types could readily be identified. To address the possibility of multiplet droplets or putative multihormonal cells, we classified individual cells using a refined set of markers highly specific to the clusters of each cell type (Methods). This cell-wise classification approach allowed for identification of ambiguous droplets that co-expressed marker gene profiles of more than one cell type. 1492 such ambiguous droplets were identified, which is on the order of the expected number of doublets in the 10X system: ∼6.4% per lane of 8000 cells, or a total of ∼2000 cells total from 4 lanes (Zheng et al., 2017). Analysis of the marker gene set combinations expressed in these cells revealed that the number of cells per combination scaled with their constituent cell population sizes. Furthermore, subpopulations of these ambiguous droplets formed separate clusters in UMAP embeddings between the main clusters with which they shared cell type markers. Taken together, these results suggested that ambiguous droplets contained more than one cell as opposed to single multihormonal cells and were removed from further analysis.

After cell filtering based on quality control metrics, 15876 cells remained for further analysis. The cell classification closely matched UMAP clusters, confirming the validity of the classification (Figure 1a). We identified six hormone producing cell types (HPC) – melanotrophs (570 cells), corticotrophs (895 cells), gonadotrophs (535 cells), thyrotrophs (133 cells), somatotrophs (4149 cells), and lactotrophs (5026 cells); two nonhormonal cell types – folliculostellate cells (FSC, 2388 cells) and pituicytes (841 cells); two vasculature cell types – endothelial cells (EC, 362 cells) and pericytes (251 cells); and blood cells – leukocytes (726 cells) and erythrocytes (164 cells, excluded from further analysis). Well known marker genes for each cell type, including hormones and hormone receptors, were highly specific to their respective cell types (Figure 1b).

**FIGURE 1.**
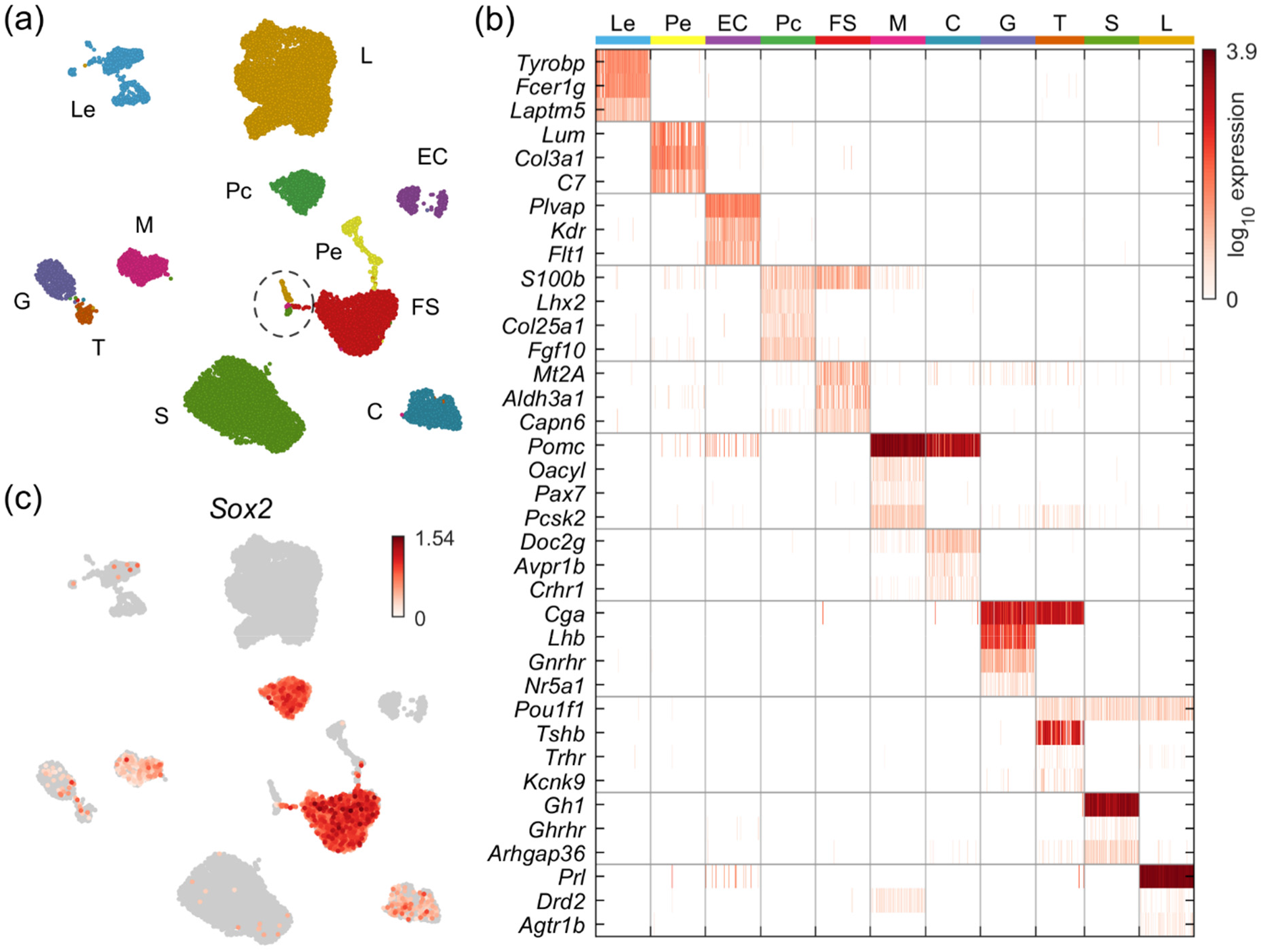
Identification of pituitary resident cell types. (a) UMAP embedding showing identified resident pituitary cell types: six hormone producing cells (HPC) – melanotrophs (M), corticotrophs (C), gonadotrophs (G), thyrotrophs (T), somatotrophs (S), and lactotrophs (L); two nonhormonal cell types (NHC) – folliculostellate (FS) cells and pituicytes (Pc); two vasculature cell types (VC) – endothelial cells (EC) and pericytes (Pe); and blood cells – leukocytes (Le). Red blood cells are excluded. Cells encircled with a dotted line express cell cycle marker genes. (b) Heatmap showing selected marker genes characteristic of identified cell types for a random subsample of up to 150 cells per type. In further analysis, we excluded Le. (c) UMAP embedding showing the expression of *Sox2*, uniformly and highly expressed among Pc and FSC with low-level expression among M, C, G, and T.

The sample identities of cells per cluster indicated the anterior lobe specificity of FSC, the posterior lobe specificity of pituicytes, and subclusters of leukocytes, pericytes, and EC that were specific to each lobe, reflecting the lobe-specific functions. A small population of cells expressing proliferation marker genes was also identified (dotted circle, Figure 1a), and cell-wise classification indicated these cells comprised primarily of *Pou1f1*-expressing lactotrophs and somatotrophs, as well as FSC, as we and others have previously observed (Cheung et al.; Fletcher et al., 2019; Ho et al., 2020; Zhang et al., 2020). Notably, however, we did not observe *Sox2* expression confined to a small cluster of putative adult pituitary stem cells. In contrast, we found that *Sox2* was a uniformly well-expressed marker gene among *S100b*-positive FSC and pituicytes, with lower-level expression in a fraction of melanotrophs, corticotrophs, gonadotrophs, and thyrotrophs (Figure 1c). This finding indicates that *Sox2* expression alone is insufficient for identification of stem cells. We therefore performed a detailed characterization of the transcriptomic signatures of FSC and pituicytes.

### 3.2 Heterogeneity of folliculostellate cells

We investigated the heterogeneity of FSC at the single-cell transcriptome level by performing UMAP embedding and unsupervised clustering of these cells. Two subpopulations were identified, one larger (FSC1, 1877 cells) and one smaller (FSC2, 511 cells) (Figure 2a). Interestingly, FSC2 cells expressed a much greater diversity of genes compared to FSC1, with a median of 3137 and 2099 genes per cell, respectively (Figure 2b), suggesting that FSC2 cells possess increased differentiation potential (Gulati et al., 2020). The FSC2 cluster also contained a small subcluster of cells expressing cell cycle marker genes (Figure 2a, circled), accounting for the same proliferative FSC seen in Figure 1a.

**FIGURE 2.**
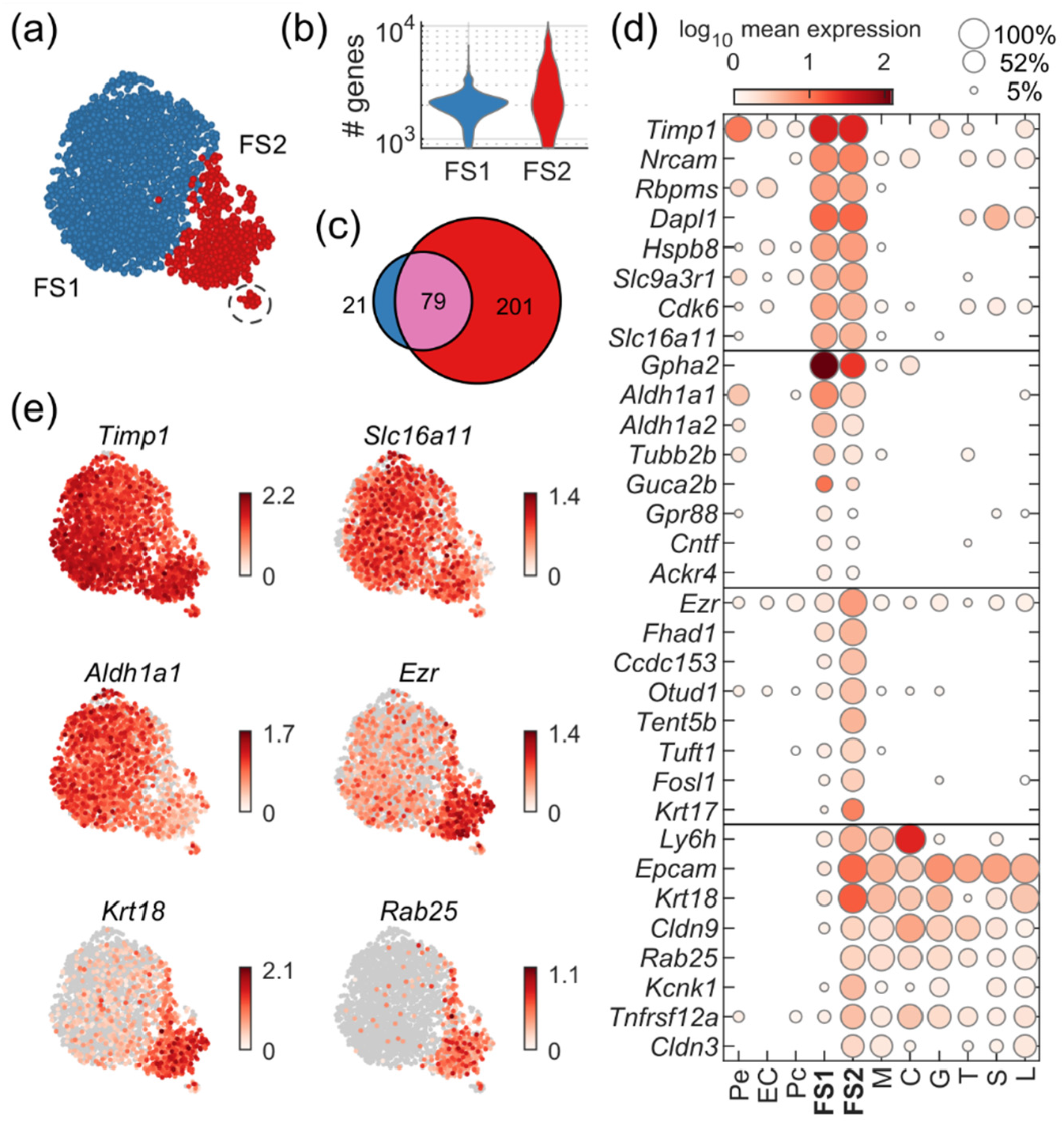
Identification of two folliculostellate cell subtypes. (a) UMAP embedding showing unsupervised clustering of FS cells into two subtypes. A small portion of FS2 cells (circled) express cell cycle markers. (b) Violin plot of the distributions of number of genes per cell in FS1 and FS2, with median values of 2099 and 3137, respectively. (c) Venn diagram showing the number of genes identified as FS1-dominant, FS2-dominant, and common to both. (d) Dot plot showing selected top FS-expressed genes: dominant in both subtypes (first group), FS1-dominant (second group), FS2-dominant (third group), and FS2 and HPC co-dominant (fourth group). (e) UMAP scatter plots showing the expression patterns of selected genes identified as dominant in both FS1 and FS2 (*Timp1, Slc16a11*), FS1-dominant (*Aldh1a1*), FS2-dominant (*Ezr*), and coexpressed by FS2 and HPC (*Krt18, Rab25*).

Pairwise differential expression analysis (Methods) was used to identify FSC1- and FSC2-upregulated genes relative to all vascular and HPC types. Pituicytes were excluded from this analysis to allow subsequent identification of shared FSC and pituicyte markers (Section 3.3). We included genes that were expressed in at least 15% of FSC subtypes, with a significant 2-fold upregulation of expression (adjusted p < 1e-6) and that had expression in at least 10% greater proportion of FSC relative to other cell types. Excluding genes upregulated also in pituicytes, this analysis identified 79 genes dominantly coexpressed by both FSC1 and FSC2 (pan-FSC), 21 FSC1-dominant genes, and 201 FSC2-dominant genes (Figure 2c). Interestingly, most FSC1-upregulated genes (79/100) were also expressed by FSC2, consistent with the closeness of these subclusters in UMAP embeddings. In contrast, many FSC2-upregulated genes were not coexpressed by FSC1, consistent with the larger diversity of expression in FSC2. These results suggest that FSC1 and FSC2 cells are highly similar, but FSC2 cells have additional activated gene expression programs.

Figure 2d highlights some of the top FSC-expressed genes, and Figure 2e shows the expression level of selected genes in individual cells, represented as the color of each point in the UMAP embedding. Notable pan-FSC genes included tissue inhibitor of metalloproteinase 1 (*Timp1*) (Azuma et al., 2015), the proton-linked monocarboxylate transporter *Slc16a11*, a marker for Müller glial cells of the retina, and *Cdk6*, important in regulation of the G1-S transition in the cell cycle. FSC1-dominant genes included glycoprotein hormone alpha-2 (*Gpha2*), co-expressed in some corticotrophs (Okada et al., 2006), and aldehyde dehydrogenase 1 family, member A1 (*Aldh1a1*) (Fujiwara et al., 2007). FSC2-dominant genes included: ezrin (*Ezr*); the OTU deubiquitinase 1 (*Otud1*), a known regulator of Hippo signaling (Yao et al., 2018); *Tent5b*, which encodes terminal nucleotidyltransferase 5B; and *Krt17*, encoding the intermediate filament keratin 17.

While investigating the extra genes expressed in FSC2, we noticed that FSC2 shared expression of several genes with HPC (Figure 2d, bottom group). Differential expression analysis identified 74 genes FSC2 upregulated relative to all other non-HPC types and also upregulated in at least one HPC cell type, while only 2 such genes were identified for FSC1. Some examples of these genes included: *Krt8* and *Krt18*, cytokeratins which have been associated with pituitary stem cell niches (Shintani & Higuchi, 2021); *Rab25*, important in apical endocytic vesicle recycling and implicated in cell migration and tumor metastasis (Caswell et al., 2007); a potassium two pore domain channel gene *Kcnk1;* the epithelial cell adhesion molecule, *Epcam*; and the tight-junction components *Cldn9* and *Cldn4*.

Taken together, the expression profiles of FSC suggest a classical FSC role for FSC1 cells, including expression of enzymes, transporters, and shared pan-FSC marker genes. In contrast, FSC2 expressed additional genes suggestive of the epithelial-like polarized cell phenotype expected for cells participating in dense parenchymal FSC clusters or marginal cell layer associated with stem cell niches. Furthermore, their shared expression of genes with FSC points to the possibility that this cluster also contains HPC-committed progenitor cells.

### 3.3 Folliculostellate cells vs pituicytes: common and cell-type-specific gene expression

Because *Sox2* was expressed in all FSC and pituicytes, we next sought to compare the transcriptomic profiles of FSC with their counterparts in the posterior lobe. Extending the gene intersection approach used for FSC, we added the set of genes upregulated in pituicytes relative to all vasculature cells and HPC. In addition to the genes identified for FSC1 and FSC2, this analysis uncovered 384 pituicyte-dominant genes, 6 pituicyte and FSC1 co-dominant genes, 33 pituicyte and FSC2 co-dominant genes, and 51 genes common to all three cell groups (Figure 3a – note the agreement of FSC1 and FSC2 exclusive regions with Figure 2c).

**FIGURE 3.**
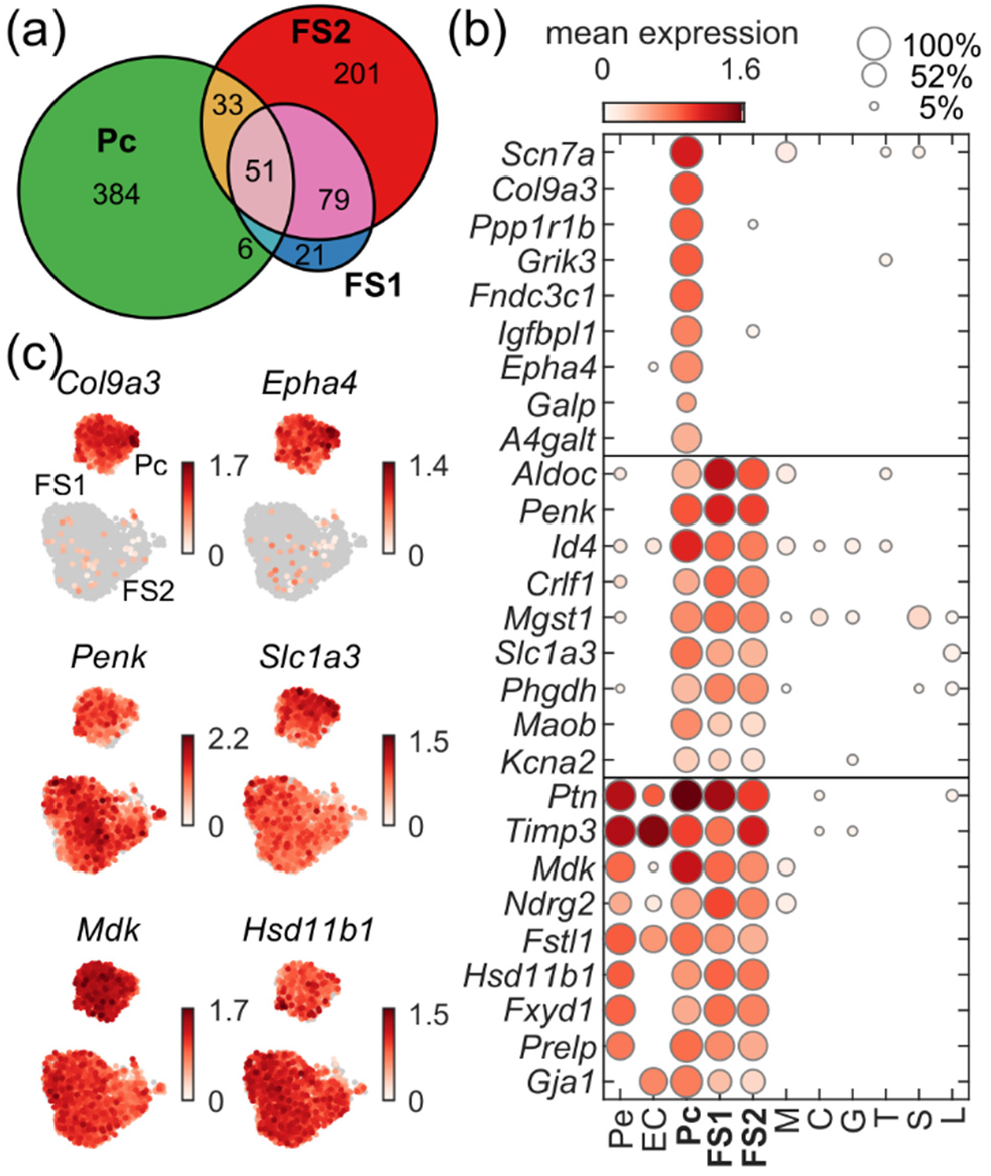
Comparison of FS and Pc gene expression profiles. (a) Euler diagram showing the number of genes identified as Pc-dominant, FS1-dominant, FS2-dominant, and common to each combination of the three cell subtypes. (b) Dot plot showing selected top expressed Pc-dominant genes (top group), genes common to both Pc and FSC (middle group), and genes coexpressed by Pc, FSC, and vascular cells (bottom group). (c) UMAP scatter plots of Pc and FSC showing the expression patterns of selected genes from panel b.

Figure 3b shows the expression profile of selected top-expressed pituicyte and FSC genes, with notable examples shown in UMAP plots in Figure 3c. Pituicytes uniquely expressed a significant number of genes (Figure 3b, top), likely reflecting the unique requirements of their function in the posterior lobe. These included ECM components such as *Col9a3*, the glial-cell expressed voltage-insensitive sodium channel *Scn7a*, and endogenous signaling genes such as galanin-like peptide (*Galp*) and ephrin type-A receptor *Epha4*.

Genes common to pituicytes and FSC (Figure 3b, middle) indicated the common identity of these cells as astrocyte-like cells. In addition to *Sox2* and *S100b* (Figure 1), these cells shared expression of astrocyte marker genes such as *Penk, Slc1a3*, and *Maob*. We also identified several genes coexpressed with pericytes and EC (Figure 3b, bottom). This comparison was motivated by examination of some genes expected to be expressed in FSC: the gap junction forming connexin 43, *Gja1*, consistent with FSC cell networks (Fauquier et al., 2001), and *Hsd11b1*, involved in cortisol negative feedback on the pituitary (Korbonits et al., 2001).

To gain a wholistic overview of the roles suggested by FSC- and pituicyte-dominant genes, we tested each of the three sets of genes for gene enrichment among GO terms (BP, Biological Process; CC, Cellular Component; MF, Molecular function) and pathways (KEGG, Kyoto Encyclopedia of Genes and Genomes; REAC, Reactome) using gProfiler (Raudvere et al., 2019). The results were then compiled to investigate shared and cell type-specific term enrichment. We investigated thematic patterns in the set of significantly enriched terms using UMAP embedding and clustering, using the overlaps among the sets of genes enriched for each term as a metric for term similarity (Methods). The resulting UMAP embedding of terms colored by cell type combinations is shown in Figure 4a, with the Euler diagram of term intersections serving as the color legend.

**FIGURE 4.**
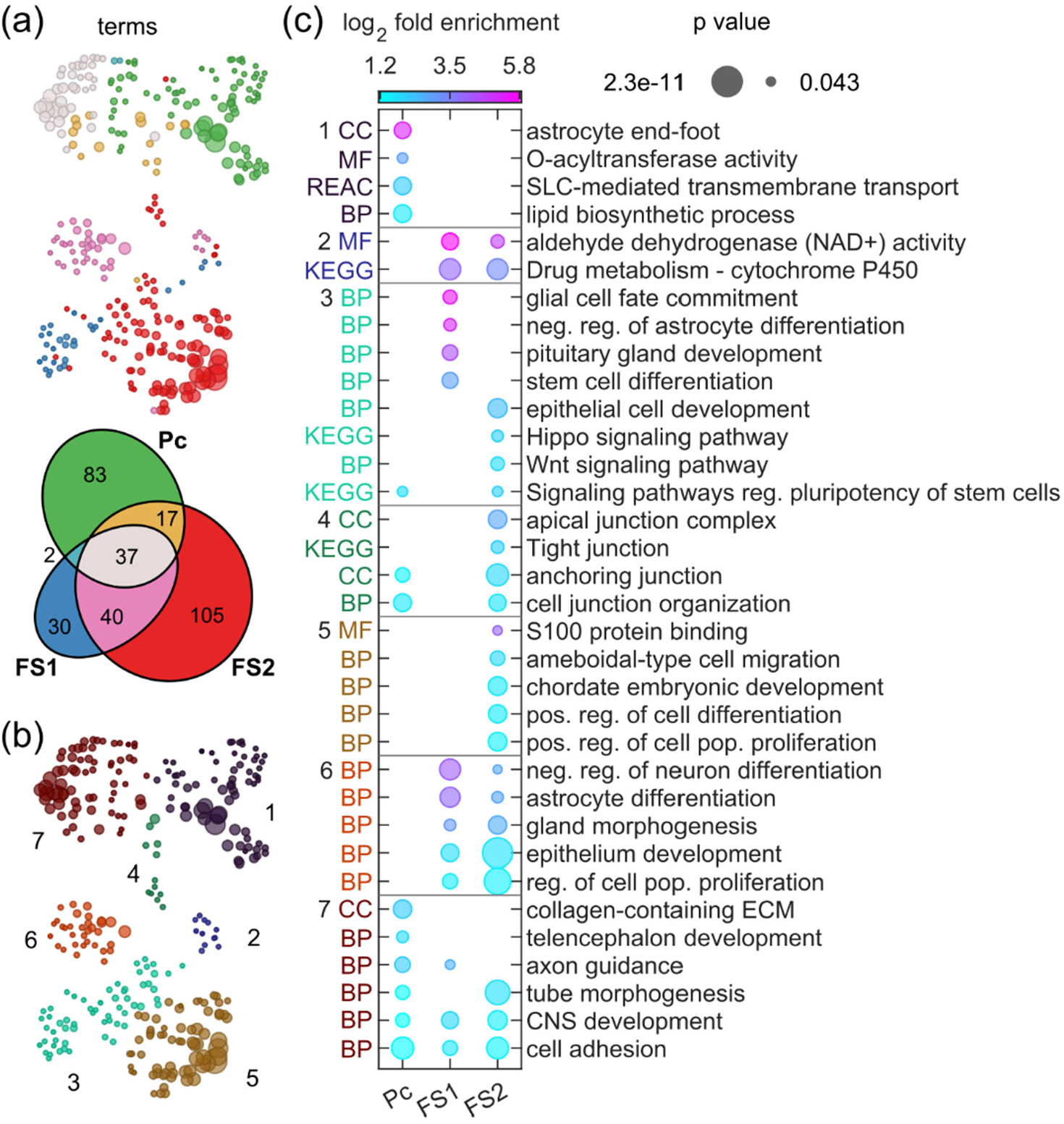
Gene Ontology and Pathway enrichment analysis of FS- and Pc-dominant genes. (a) UMAP plot of GO and pathway enriched terms for FSC and Pc. The Euler diagram indicates the number of terms enriched for each combinations of cell types. (b) Clustering of terms to identify patterns. (c) Dot plot indicated the fold-enrichment (color) and p-value (size) for selected terms per cluster.

Clustering of terms (Figure 4b) was used to group together terms with similar sets of enriched genes, and the enrichment test results for representatives from each cluster are shown in Figure 4c. Clustering revealed groupings of terms suggesting several overlapping and distinct functions for pituicytes and FSC, including metabolism and detoxification, differentiation/development/ morphogenesis, signaling, and ECM and cell junctions. Clusters 1 and 2 point to the functions of pituicytes and FSC relating to metabolic processes and detoxification, as well as multiple types of metabolite and ion transporters in pituicytes.

Many terms arise for developmental processes and control of cell proliferation (clusters 3-7). Cluster 3 points to roles for FSC1 in promoting differentiation of cells, while FSC2 genes were enriched instead in related signaling pathways involved in regulation of stem cell potency, such as Hippo and Wnt, as well as epithelial-like cell development. Cluster 4 captured terms related to FSC2-mediated tight junctions, consistent with formation of dense FSC clusters and suggestive of involvement in stem cell niche formation by these cells. Cluster 5 contained FSC2-enriched terms related to cell migration, development, and positive regulation of cell proliferation, further suggestive of stem cell-like activity. Cluster 6 featured terms enriched for both FSC1 and FSC2 genes, pointing to shared involvement in the regulation of differentiation of astrocyte-like cells and “neuron-like” HPC, gland morphogenesis, and general regulation of cell proliferation. Finally, cluster 7 pointed to pituicyte-specific roles involving collagen-containing ECM, development processes unique to the posterior lobe, and more generic terms relating to CNS development, axon guidance, and cell adhesion.

### 3.4 Development and differentiation marker gene expression pituitary cell types

We next investigated the expression of additional reported stem/progenitor and differentiation markers, as well as core signaling pathways. Figure 5a shows a dotplot including a selection of these genes, grouped by expression pattern, and Figure 5b shows UMAP embeddings colored by the expression per cell of six key genes. Markers including *S100b, Sox2*, and *Sox9* were coexpressed specifically in pituicytes and FSC, while *Vim, Cd9*, and *Hes1* were coexpressed additionally in pericytes and EC (Figure 5a, top group). Pituicytes uniquely expressed many genes, including expected markers *Fgf10, Tbx3, Lhx2*, and *Nkx2-1*, reflecting differences in developmental/differentiation processes in the posterior lobe. FSC specifically expressed *Prop1, Hey1, Hey2*, and *Msx1*, and shared expression of *Prrx1* and *Heyl* with pericytes. They also coexpressed several differentiation marker genes with HPC, such as *Isl1, Cd24, Six1, Six6, Pitx1, Pitx2*, and *Lhx3*, complementing the FSC2-HPC shared genes shown in Figure 2.

**FIGURE 5.**
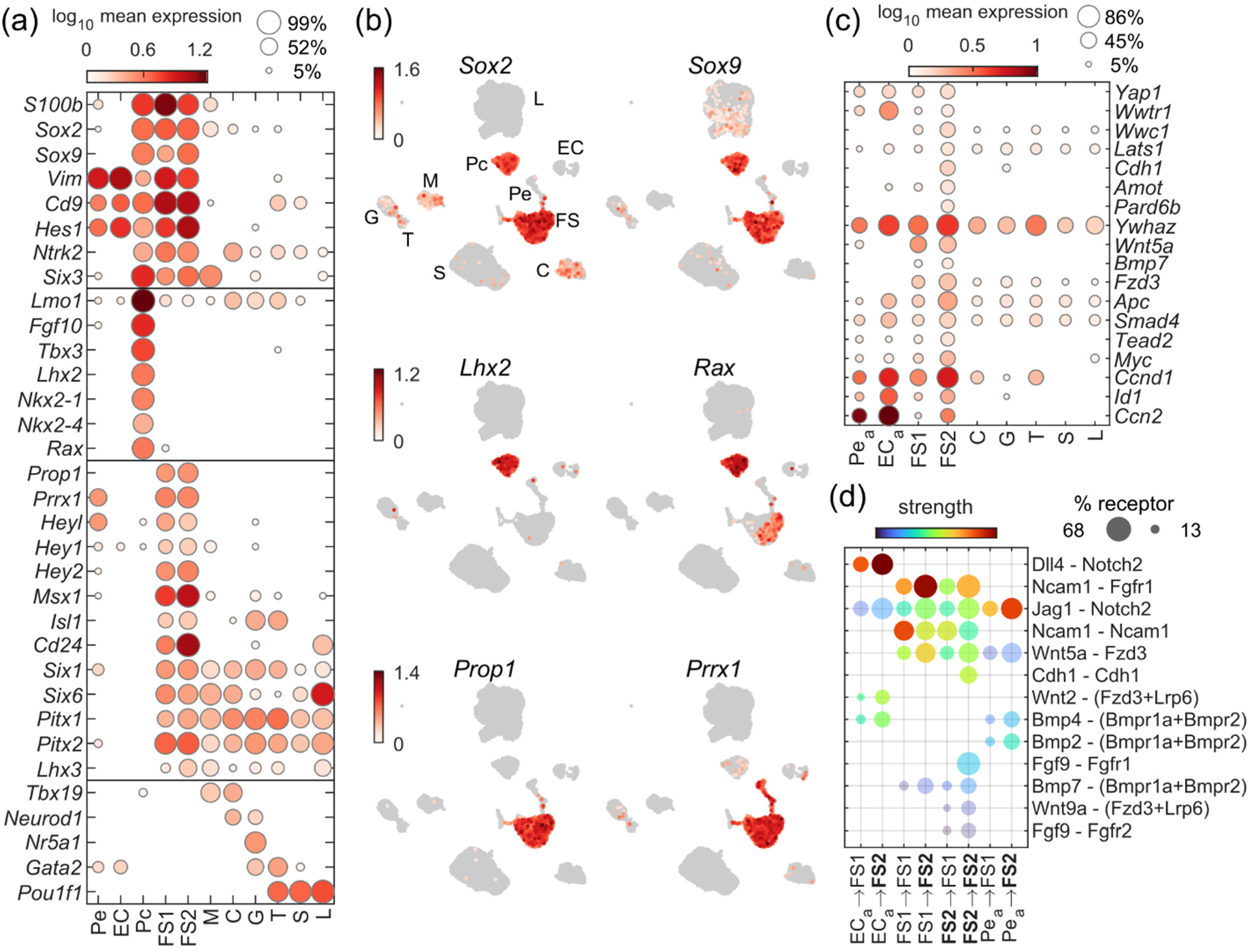
Development and differentiation signature genes. (a) Dotplot showing the mean expression and percentage of cells expressing selected genes representative of developmental and differentiation processes in the pituitary. (b) UMAP embeddings showing the expression per cell for selected marker genes. (c) Dot plot showing anterior lobe expression of key genes from the KEGG Hippo signaling pathway term. (d) FSC-targeted ligand-receptor (L-R) interactions from core signaling pathways based on CellChat analysis (Methods). Rows represent L-R interactions, and columns labels indicate source-target directionality. Point color represents potential interaction strength and size indicates the percentage of target cells expressing the receptor (or the minimum % expression among receptor complex genes). All points shown have bootstrap p-value < 1e-4, and both source and target cell types express the ligand or receptor gene in at least 10% of cells.

Motivated by the enrichment of FSC2 genes in Hippo and Wnt signaling pathway terms (cluster 3, Figure 4), we examined expression of all genes from the KEGG Hippo signaling pathway term (KEGG:rno04390), focusing on FSC of the anterior lobe. 90 pathway genes had expression in at least 15% of either FSC1 or FSC2. However, the proportion of cells expressing these genes was over 10% greater in FSC2 than in FSC1 for 81 of these, suggesting a higher degree of Hippo signaling activity in FSC2. We highlight the anterior-lobe expression profiles of key pathway genes in Figure 5c.

Both *Yap1* and Taz (*Wwtr1*), the main coactivators of the pathway, were better expressed in FSC2 compared to FSC1. FSC2 also expressed genes coding for the transcription enhancer factor (TEAD/TEF) family of transcriptional factors (*Tead2* and *Tcf7l1*). While *Sox2* was expressed uniformly among FSC, several other downstream Yap1 activated target genes were upregulated in FSC2 compared to FSC1, including *Ccnd1, Myc, Ccn2, Id1/2*, and *Nkd1*.

FSC2 also expressed genes suggesting regulation of Hippo by cell density in these cells, consistent with our results indicating these cells likely participate in cell junctions needed for formation of dense FSC clusters or the marginal cell layer. including cell polarity regulator *Pard6b, Amot, Amotl2*, and E-cadherin (*Cdh1*), the latter also noted as a pituitary stem-cell marker gene. Several Hippo-regulating signaling pathway genes were also well expressed. *Wnt5a* ligands were more highly expressed by FSC1, while FSC2 expressed Fzd receptors at higher levels, suggesting FSC1 to FSC2 signaling. BMP signaling pathway elements were more highly expressed in FSC2, including *Bmp7* and Bmp receptors, suggesting possible autocrine BMP signaling.

We tested the possibility of FSC-targeted Wnt, Bmp, Notch, and FGF signaling pathways, as well as cell-cell interactions, using the CellChat approach for estimating ligand-receptor interactions (Jin et al., 2021). Selected results are shown in the dot plot in Figure 5d, which indicates potential strength of interaction (color) as well as the percentage of cells of the target cell type that expressed the receptor (size). We note that this analysis shows only the possibility for interaction – in addition to gene expression, the cells would need to produce functional proteins, and would need to be physically oriented relative to each other in a way that permits the interaction. Among the strongest interactions included Notch2 signaling via EC-expressed Dll4 ligands or pericyte-expressed Jag1 ligands. Also identified were strong homotypic Ncam1 interactions among FSC1, as well as Ncam1-Fgfr1 interactions directed from FSC1 to FSC2. The strongest interaction among Hippo regulating pathways involved FSC1 to FSC2 signaling via *Wnt5a* and *Fzd1* or *Fzd3* receptors (the two most highly expressed Fzd receptors, among others), as well as homotypic FSC2 *Cdh1* cell-cell interactions. Further Wnt signaling may be mediated by EC-expressed *Wnt2*. BMP signaling potential was strongest via EC-expressed *Bmp4* and pericyte-expressed *Bmp2* directed towards FSC2, followed by the possibility of *Bmp7* autocrine signaling in FSC2 cells. Finally, we also identified a possible autocrine signaling of *Fgf9* in FSC2 via the *Fgfr1* or *Fgfr2*. Taken together, these results indicate that FSC2 cells are likely members of stem cell niches, and active as stem cells or HPC-committed progenitors.

### 3.5 Expression of S100 family members

S100b, a classical glial cell marker, has historically been viewed as a definitive marker for FSC in the pituitary. More recently, evidence for the expression of various S100 gene family members has emerged in the pituitary. We confirmed the expression of several S100 gene family members in vascular and nonhormonal cell types. *S100b* was a good marker for FSC and pituicytes, although some low-level expression in some M and pericytes was also observed. *S100a1* had a similar expression pattern but was better expressed in FSC than pituicytes (Figure 6a, b). *S100a6* and *S100a11* were highly specific to FSC and pericytes, but not pituicytes, with stronger expression in anterior lobe pericytes (Pe_a_). *S100a10* was similarly well expressed in FSC and pericytes, but also exhibited lower-level expression among EC, and *S100a10* and *S100a11* had some low-level expression in HPC. *S100a13* was broadly expressed, but with highest levels in FSC and pericytes, while *S100a16* was localized to all non-HPC, predominantly EC and FSC. Finally, *S100g* had a unique patter of expression, specifically localized to pituicytes of the posterior lobe and L in the anterior lobe, with low levels detected in some S and G.

**FIGURE 6.**
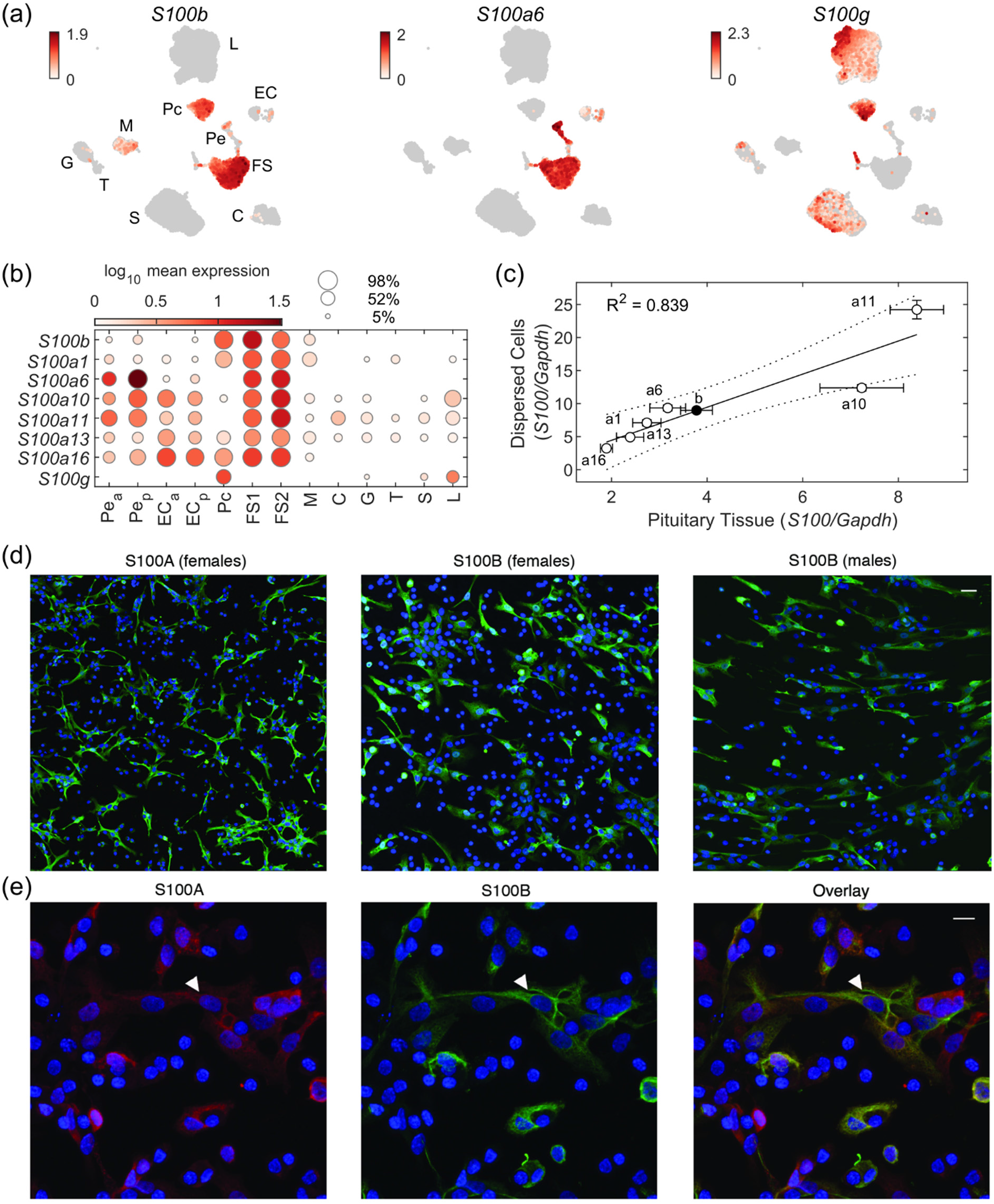
S100 gene expression in the pituitary gland. (a) UMAP scatter plots showing expression of *S100b*, common to FS and Pc, *S100a6*, expressed in FS and Pe, and *S100g*, expressed in Pc and a subset of S and L. (b) Dot plot indicating the mean expression of S100 genes in each cell type, with EC and Pe divided by lobe of origin and FSC divided by subtypes. (c) qRT-PCR measurements of S100 gene expression in dispersed cells and pituitary tissue from 44-day old female rats. Solid line – linear fit; dotted lines – 95% confidence intervals. (d, e) Immunofluorescence analysis of S100A and S100B proteins in primary cultured pituitary cells. (d) S100A expression in cells from females (left) and S100B expression in cells from females (center) and males (right). (e) Colocalization of S100A (red, left) and S100B (green, center), with overlay showing dual labeled cells (right). In 6 separate replicates, a total of 494 cells were S100A+, 339 were S100B+, and 275 were dual labeled. Cell nuclei are stained with DAPI (blue). Arrows indicate an example cell that co-expresses both proteins. Scale bars (applies to all images), 10 µm.

To confirm the expression of these S100 family members in the pituitary, we performed independent qRT-PCR and immunofluorescence analysis. qRT-PCR showed good agreement between dispersed and intact tissue samples for S100B and the six S100A family members described above (R^2^=0.839), indicating that cell dispersion did not disrupt expression of these genes (Figure 6c). In primary cultured pituitary cells from both female and male animals, S100A and S100B immunofluorescence was localized to cells with prominent projections, characteristic of FSC (Figure 6d). Colocalization of S100A and S100B occurred in 56% of S100A+ cells and 81% of S100B+ cells, counted from a total of 6 separate replicates (Figure 6e), consistent with the scRNAseq result that cell populations other than FSC, including pericytes, expressed S100A gene family members but not *S100b*.

### 3.6 Astrocyte signatures of folliculostellate cells and pituicytes

A major theme in the results described above is that FSC and pituicytes share expression of many astrocyte/glia marker genes. We finalize our analysis of FSC and pituicytes marker genes, the gene scores for which are shown in Figure 7a (FSC, top; pituicytes, bottom), by comparing them to some published marker gene lists for similar cell types. Figure 7b shows the overlap between FSC1 or FSC2 markers or pituicytes markers with published marker gene sets, including astrocytes (Batiuk et al., 2020; Lovatt & Nedergaard, 2012), mouse pituicytes (Q. Chen et al., 2020), tanycytes (Campbell et al., 2017), and the PangolaDB marker sets for astrocytes (Franzen et al., 2019). While both cell types show similar overlap with astrocyte marker gene lists, pituicytes show a higher degree of overlap with the pituicyte and tanycyte marker lists.

**FIGURE 7.**
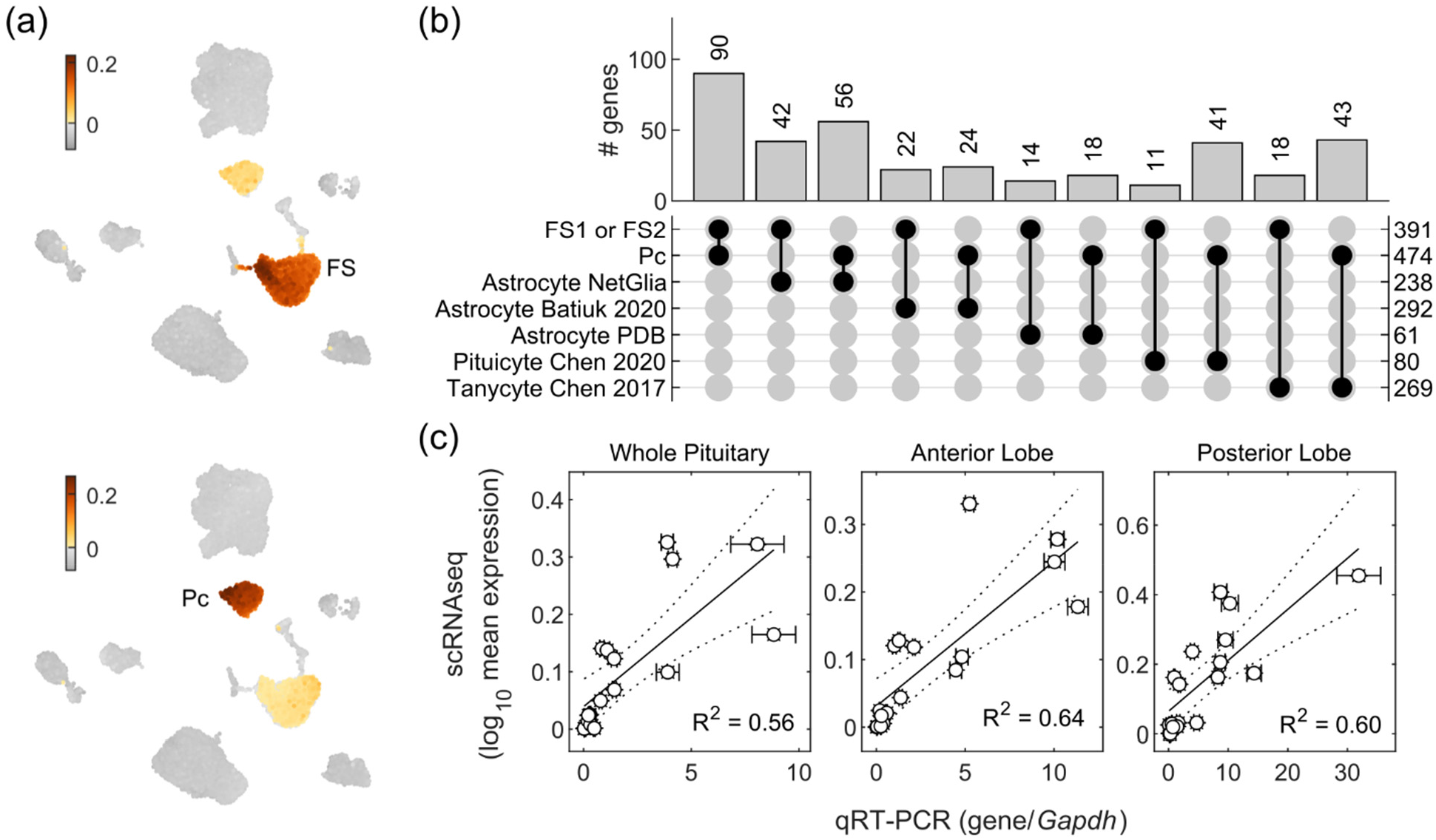
Astrocyte-like marker gene signature of FS and Pc cells. (a) Gene scores based on the list of dominant genes identified for FS1 and FS2 (top panel) and Pc (bottom panel). (b) Upset plot showing the number of common genes between FSC and Pc gene sets and published lists of marker genes for astrocytes, pituicytes, and tanycytes. (c) Correlation of the mean expression of all cells in each scRNAseq sample with qRT-PCR expression for 18 astrocyte marker genes.

To validate the presence of astrocyte marker genes, we performed qRT-PCR for 18 such genes (Figure 7c). The qRT-PCR expression levels correlated well with the mean expression of the same genes in each sample of the scRNAseq data, validating our scRNAseq findings. Genes tested included *S100b, Sox2, Gfap, Aldh1a1, Aldh1l1, Aqp4, Edn3, Fgfr3, Fibin, Gstm7, Maob, Nes, Penk, Slc1a2, Slc1a3, Sox9, Sult1a1*, and *Vim*.

## 4 DISCUSSION

We used here scRNAseq, qRT-PCR, and immunohystochemistry to further characterize the transcriptome profiles of the anterior and posterior pituitary cells obtained from adult female rats. We identified six HPC: melanotrophs, corticotrophs, gonadotrophs, thyrotrophs, somatotrophs, and lactotrophs; two nonhormonal cell types: FSC and pituicytes; and two vascular cell types: EC and pericytes. A small population of cells expressing proliferation marker genes was also identified. Separate scRNAseq analysis of anterior and posterior lobe tissue served as an internal control to confirm the anterior lobe specificity of FSC, the posterior lobe specificity of pituicytes, the predominant separation of melanotrophs of the intermediate lobe with the posterior lobe, and lobe-specific clusters of vascular cells. The anterior pituitary cell identification in this study was highly comparable with our previous work (Fletcher et al., 2019), but the percentage of specific pituitary cell population was somewhat different, probably reflecting the impact of cell dispersion procedures. The transcriptome profile of rat pituicytes described here is comparable to that reported in mouse pituicytes (Q. Chen et al., 2020) but we provide additional information about their transcriptomes and functional relationship with FSC. However, our cluster analysis did not lead to identification of a separate adult pituitary stem cell cluster.

The first scRNAseq study of whole mouse pituitary cells identified a *Sox2*-positive cluster as stem cells but did not consider FSC in their cell type classification. The transcriptome profile reported for that cluster did not list other known stem cells genes, such as *Cdh1, Cd9, Lhx3, Prop1, Prrx1, Vim, Sox9*, and *S100b* (Garcia-Lavandeira et al., 2012; Gleiberman et al., 2008; Horiguchi et al., 2018; Rizzoti, 2010; Vankelecom & Chen, 2014), but includes *Aldh1a2, Aldh3a1, Cdh26, Cpxm2*, and *Rbpms*, in our study expressed in FSC (Cheung et al., 2018). It is known, however that *S100b* is a poor marker in mouse pituitary, and recently *Aldoc* has been suggested as a good marker for FSC in both mouse and rat (Fujiwara et al., 2020). Another mouse pituitary scRNAseq study also labelled Sox2-positive cells as stem cells and did not attempt to identify FSC (Lopez et al., 2021). Another group separated *Sox2*-positive from FSC (Ho et al., 2020). However, their *Sox2*-positive cells co-express *Aldoc, Crlf1, Phgdh, Nrcam*, and *Mgst1*, common FSC genes, while their FSC expressed *Pdgfrb, Dcn, Lum, Col1a1* and *Col3a1*, which in our study were markers of pericytes. However, no pericyte clusters were identified in their study, which could explain this difference. A recent study with mouse pituitaries acknowledged that stem cells were likely a subset of FSC but classified all such cells as two stem cell clusters (Vennekens et al., 2021).

The expression of *Sox2* in FSC is consistent with the old idea that FSC have stem cells capability (Inoue et al., 2002). However, renaming FSC as *Sox2* positive stem cells seems unreasonable, as these cells are an established anterior pituitary cell lineage with numerous specific functions reported during a long history of studies (Le Tissier & Mollard, 2021). These include their capacity for generation of numerous endogenous ligands, detoxification enzymes, extracellular matrix proteins, and cell adhesion molecules (Fletcher et al., 2019), which are critical for the proper tridimensional organization and function of the anterior pituitary gland, which includes the FSC gap-junction coupled network (Fauquier et al., 2001). We show here that *Sox2*, the putative stem cell marker gene of the anterior pituitary, is expressed not only by FSC, but also by pituicytes and, at lower levels, by some melanotrophs, corticotrophs, gonadotrophs, and thyrotrophs, consistent with our previous experiments done with anterior pituitary cells from adult male and female rat (Fletcher et al., 2019). Therefore, the heterogeneity of *Sox2*-positive cells is obvious and cannot be related to clustering problems because our study also included pituicytes and melanotrophs from the posterior pituitary preparation. Such heterogeneity also clearly indicates that *Sox2* cannot be accepted as a sole marker gene for stem cells, motivating the need for a more comprehensive set of markers.

This prompted us to investigate heterogeneity of FSC to address the hypothesis that a fraction of FSC act as stem cells. Our analysis suggested that FSC form at least two subclusters. FSC1 contained more cells (_∼_80%) but less diverse genes, as opposed to FSC2 with fewer cells (_∼_20%) expressing very diverse genes. Several findings indicate that FSC2 may represent active stem/progenitor cells in anterior pituitary. Hippo signaling has been shown to be important in pituitary stem cell maintenance (Lodge et al., 2016), and a large complement of these genes were more highly expressed in FSC2. Proliferative FSC were clustered here with FSC2 and proliferative FSC were observed in our previous scRNAseq data (Fletcher et al., 2019), suggestive of stem cell self-renewal and/or asymmetric cell division required to generate HPC-committed cells. Several Gene Ontology terms were FSC2-specific for development, including positive regulation of cell proliferation, and stem cell pluripotency regulation. FSC2 specifically expressed the genes coding for tight junction proteins, consistent with the idea that these cells were members of the parenchymal and/or marginal zone stem cell niches (Higashi et al., 2021). This analysis also Identified FSC2-specific terms for cell migration, which could be related to the epithelial-mesenchymal transition needed for release HPC-committed cells from stem cell niches (S. Yoshida et al., 2016). Our analysis also revealed that FSC2 shared genes with HPC, including *Rab25*, which encodes low molecular weight GTPase with preferential expression in the pituitary gland (St-Amand et al., 2011) and has been linked to cell migration in tumor metastasis (Caswell et al., 2007), keratins encoded by *Ktr8, Krt18*, and a potassium two pore domain channel gene *Kcnk1*, as well as *Epcam*, which encodes the epithelial cell adhesion molecule expressed in nonfunctional pituitary adenomas (Wang et al., 2019). Such shared expression profiles are suggestive of cells in transition to HPC.

To better understand the nature and roles of nonhormonal cells, we compared the expression of dominant genes in FSC1, FSC2, and pituicytes. Among the dominant genes, only _∼_12% are common for three clusters, indicating a common function of these cells in pituitary functions. These include the expression of receptor ligand genes, stem-like genes, and astroglial-like genes. Receptor ligand genes include the astrocyte marker genes *Mdk* (Y. Yoshida et al., 2014) and *Ptn* (Yeh et al., 1998), whose proteins act as signaling molecules during pituitary development (Fujiwara et al., 2014). *Penk*, another gene expressed in astroglia (Theodoridu et al., 1994), encodes proenkephalin, a stable surrogate for enkephalins, endogenous opioid peptides. Opioid receptor genes are not expressed in HPC, FSC, and pituicytes, but receptors are detected in the posterior pituitary gland (Stojilkovic et al., 1987), indicating their expression in the terminals of oxytocin and arginine vasopressin-secreting neurons. However, the opioid growth factor receptor gene *Ogfrl1* is expressed in the fraction of FSC and HPC, as well as in pituicytes and EC.

In addition to *Sox2*, FSC1, FSC2, and pituicytes expressed other anterior pituitary stem cell marker genes, including *Sox9, Cd9, Vim, Hes1, Six3, Ntrk2*, and *Gpr37l1*, among others. This is consistent with a hypothesis that FSC and pituicytes have similar identities, and that pituicytes or a fraction of these cells may have a stem cell-capacity analogous to FSC2 in the anterior pituitary. However, other developmental/differentiation genes were expressed in a cell-type-specific manner; FSC1 and FSC2 cells expressed *Prop1, Prrx1, Heyl, Hey1, Hey2, Msx1, Isl1, Cd24, Six1, Pitx1, Pitx2*, and *Lhx3*, while pituicytes expressed *Lmo1, Fgf10, Tbx3, Lhx2, Nkx2-1, Nkx2*-*4*, and *Rax*. It is also important to stress that *Sox2* and *Sox9* are astrocyte marker genes (Xia & Zhu, 2015), as well as *Vim* (Kamphuis et al., 2015), *Hes1* (Imayoshi et al., 2013), *Six3* (Yu et al., 2020), *Ntrk2* (Saba et al., 2018), and *Gpr37l1* (La Sala et al., 2020).

*S100b* is also a good marker for FSC and brain astrocytes (Osuna et al., 2012), as well as for pituicytes (Q. Chen et al., 2020). *S100b*-positive pituicytes were reported to appear in the posterior lobe on rat embryonic day 15.5 and in the anterior lobe on embryonic day 21.5 (Horiguchi et al., 2016). S100b expressing FSC were shown to have capacity to differentiate into HPC in vivo and in vitro (Higuchi et al., 2014). Like *Sox2*, we also detected *S100b* in a fraction of rat melanotrophs and corticotrophs. S100 proteins are a multigenetic family of low molecular weight calcium binding proteins, consisting of several related groups of molecules, in addition to S100b: multiple S100a proteins, S100g, S100p, and S100z (Donato et al., 2013). Several reports indicate S100g proteins in lactotrophs and GH3 immortalized lacto-somatotrophs (Nguyen et al., 2005; Vo et al., 2012). Here we confirmed that *S100g* is expressed in lactotrophs, as well as in pituicytes and a fraction of somatotrophs and gonadotrophs, but not in FSC1 and FSC2. S100a gene expression has not been previously assessed in the anterior and posterior pituitary gland. Our data indicate that FSC and pituicytes express *S100a1* and *S100a16*, the latter being also expressed in EC and pericytes. *S100a6, S100a10* and *S100a11* are expressed in FSC and pericytes, but not in pituicytes. The role of these genes in pituitary cell functions is unclear. In general, S100 proteins act as intracellular regulators, similar to calmodulin and troponin-C, but also as extracellular signaling proteins for regulating target cell activities in a paracrine and autocrine manner (Donato et al., 2013).

Pituicytes are a generally accepted subtype of astrocytes (Verkhratsky & Nedergaard, 2018), while FSC have been recognized as astrocyte-like cells (Osuna et al., 2012). As mentioned above, both cell types expressed several genes present in brain astrocytes: *Mdk, Ptn, Penk, Sox2, Sox9, Cd9, Vim, Hes1, Six3, Ntrk2, Gpr37l1*, and *S100b*, as well as *Aldoc, Aqp4, Gja1, Maob, Phgdh*, and *Slc1a3*, additional astrocyte marker genes (Verkhratsky & Nedergaard, 2018). FSC and pituicytes also expressed additional genes identified in brain astrocytes: *C1r, Crym, Gstm1, Cybrd1, Gas, Gpm6b, Id4, Lcn2, Metrn, Mxra8, Nqo1, Pcdh7, Plat, Slc20a2, Slit2, Sod3, Ttyh1, Tgfb2, Wnt5a*, and *Fzd1*. FSC, but not pituicytes, additionally expressed: *Aldh1a1, Aldh1a2, Aldh3a1, Anxa1, Cryab, Cxcl1, Fibin, Hey2, Hspob8, Il33, Msx1*, and *Slc6a8*. Finally, we observed low levels of *Gfap* expression uniquely in pituicytes.

Four genes present in mouse and rat pituicytes, *Adm, Col25a1, Rax*, and *Scn7a*, are also considered markers of hypothalamic tanycytes (R. Chen et al., 2017; Miranda-Angulo et al., 2014), but rat pituicytes and FSC do not express *Sprr1a*, a new marker gene for tanycytes (Campbell et al., 2017). Others have reported that tanycyte-specific genes also include *Col23a1, Frzb, Lhx2, Ptn, Slc16a2, Slc17a8, and Vcan* (R. Chen et al., 2017). Rat pituicytes expressed *Lhx2*, but not *Col23a1, Frzb*, and *Slc17a8*, both pituicytes and FSC express *Slc16a* and *Ptn*, and *Vcan* is expressed by FSC only. Comparison of rat FSC and pituicyte transcriptome profiles with mouse whole brain astrocytes shows that all three cell types express *Slc1a3*, an astrocytic glutamate transporter, as well as *Atp1b2, Glul, Aqp4, Fgfr3, Ogt, Fam107a, Sox9, Apoe, Gabrg1, Tryl*, and *Tlr3;* FSC and brain astrocytes express *Id3, Agt*, and *Chrdl1*; both pituicytes and brain astrocytes expressed *Mertk* and *Mfsd2a* (Batiuk et al., 2020). We also compared rat pituicytes and FSC with other scRNAseq analysis in terms of astrocyte scores and found that both cell types satisfied this criterion and pointed to quantitative rather than qualitative differences between FSC and pituicytes.

In conclusion, our experiments clearly show that *Sox2* is not a specific marker gene of active stem cells in pituitary gland, but a gene expressed uniformly among FSC1, FSC2, and pituicytes. These cells also widely expressed several other reported pituitary stem cell marker genes, including *S100b, Sox9, Cd9, Hes1*, and *Six3*, consistent with the hypothesis that they have stem/progenitor capacity. In the anterior pituitary, FSC1 lacked expression of genes consistent with stem cell niche or active stem cell state, and likely correspond to “classic” FSC. In contrast, FSC2 expressed genes consistent with stem cell activity and niche formation, likely consisting of active stem cells and HPC-committed progenitors. The finding that *Sox2* and *S100b* are expressed in fractions of HPC, and *Lhx3* in most of them, suggests that the transition from stem/progenitor to differentiated cells continues in adult animals. As HPC age, *Sox2*/*S100b* expression may decrease, unlike *Lhx3*, which is required for normal function of these cells. A higher percentage of S*ox2*-positive melanotrophs and corticotrophs than other HPC may indicate a shorter half-life of *Tbx19*-expressing cells. On the other hand, pituicytes have an additional set of development/differentiation genes and are physically separated from the anterior lobe, which provides a mechanism for cell renewal independently of the anterior pituitary. Finally, the transcriptome profiles of FSC and pituicytes indicates that both cell types met the criteria for astroglial cells, which deviate significantly from hypothalamic astrocytes.

## ACKNOWLEDGEMENTS

This study utilized the high-performance computational capabilities of the Biowulf Linux cluster at the National Institutes of Health, Bethesda, MD (http://biowulf.nih.gov). This work was supported by National Institutes of Health grants from the Intramural Research Program of the Eunice Kennedy Shriver NICHD and NIDDK.

